# Beware of asymptomatic transmission: Study on 2019-nCoV prevention and control measures based on extended SEIR model

**DOI:** 10.1101/2020.01.28.923169

**Authors:** Peng Shao, Yingji Shan

## Abstract

**Background:** The 2019 new coronavirus, “2019-nCoV”, was discovered from Wuhan Viral Pneumonia cases in December 2019, and was named by the World Health Organization on January 12, 2020. In the early stage, people knows little about the 2019-nCoV virus was not clear, and the spread period was encountering China’s annual spring migration, which made the epidemic spread rapidly from Wuhan to almost all provinces in China.

**Methods:** This study builds a SEIRD model that considers the movement of people across regions, revealing the effects of three measures on controlling the spread of the epidemic.Based on MATLAB R2017a, computational experiments were performed to simulate the epidemic prevention and control measures.

**Findings:** The research results show that current prevention and control measures in China are very necessary. This study further validates the concerns of international and domestic experts regarding asymptomatic transmission (E-status).

**Interpretation:** The results of this study are applicable to explore the impact of the implementation of relevant measures on the prevention and control of epidemic spread, and to identify key individuals that may exist during the spread of the epidemic.

## 1 Introduction

The 2019 new coronavirus, “2019-nCoV”, was discovered from Wuhan Viral Pneumonia cases in December 2019, and was named by the World Health Organization on January 12, 2020. In the early stage, people knows little about the 2019-nCoV virus was not clear, and the spread period was encountering China’s annual spring migration, which made the epidemic spread rapidly from Wuhan to almost all provinces in China. As of 20:30 on January 27, 2020, 2844 cases were confirmed nationwide, 5794 were suspected, 58 were cured, and 81 died. Such a major epidemic is a serious challenge to people’s lives and an important test of public health emergency management capabilities.

In recent days, domestic and foreign scholars have published research results about new coronaviruses online, and some of them are non-medical scholars who use mathematics and computer technology to simulate and predict disease transmission. Scholars from the field of public health present a timely evaluation of the Chinese 2019-nCov epidemic in its initial phase[1]. Different from the existing studies, this study taking into account the new characteristics of new coronaviruses and the current epidemic prevention and control measures in China, and constructing a SEIRD model that considers the movement of people across regions. The purpose of this study is to reveal the role of the three most important current measures to control the spread of the epidemic, such as quarantine of infected persons, reduction of human mobility, and improvement of treatment.

## 2 Model

The warehouse model was proposed by Kermack and McKendrick in 1927. According to the characteristics of actual transmission behavior, there can be multiple states of an individual. The two most basic states are Susceptible (S) and Infected (Infected, I). According to the types of individual states included in the model, classic warehouse models such as SI model[2], SIS model[3], SIR model[4], and SEIR model [5]. The SI model is a basic model, and other warehouse models are derived models built according to research needs. The SEIR model considers Exposed, that is, vulnerable individuals are infected, but they cannot infect other vulnerable individuals within a certain incubation period.

On the basis of the SIR model, considering that 2019-nCoV is an infectious disease with a latent period, the E (Exposed) state and D (Death) state are added to construct the SEIRD propagation model. The schematic diagram of the SEIRD propagation model is shown in figure 1.

**Figure 1.**
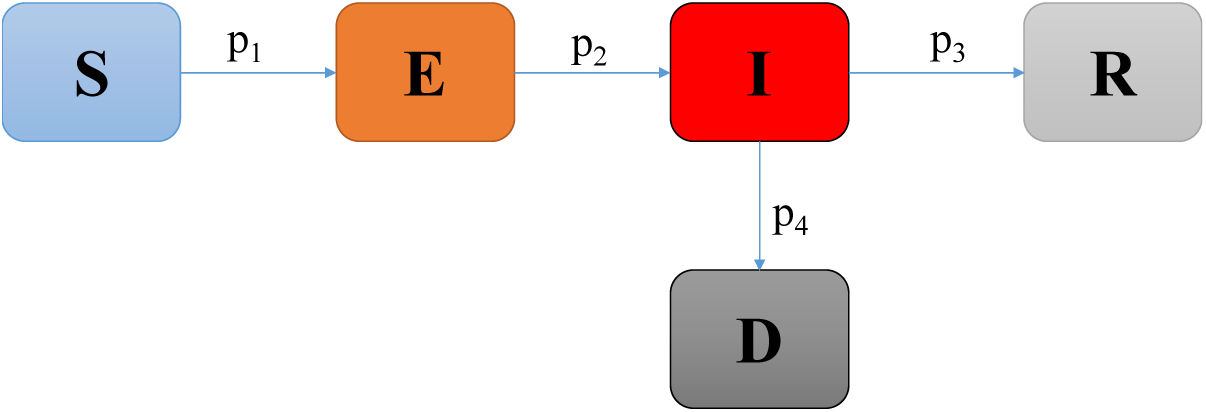
SEIRD model.

The members in this model mainly have the following four conversion methods.

- First, S → E. Relevant evidence shows that during the 2019-nCoV infection process, not only confirmed patients have infectious capacity, but also those asymptomatic transmission individuals also have the ability to infect others. Therefore, in the SEIRD model, the susceptible status (S) will change to a Exposed status (E) with a certain probability after contacting the infected individual (I) or the Exposed individual (E).
- Second, E → I. Relevant evidence shows that the longest incubation period for 2019-nCoV is 14 days and the shortest is 1 day. The Exposed status (E) may be transformed into an infected status (I) after the incubation period.
- Third, I → R. The infected individual (I) will be isolated and treated with a certain probability in hospital and will change to the R status.
- Fourth, I → D. Relevant evidence shows that infected patients die after 15 days without effective treatment.

During the 2019-nCoV transmission, the movement of individuals in the I and E status caused a rapid spread of the epidemic. Therefore, during the spread of the epidemic, consideration should be given to the contagion caused by individuals moving in different regions / communities / urbans. Suppose there are NU members in a two-dimensional space. In the initial stage, NU members are randomly and uniformly distributed in NC * NC communities (also can be understood as cities), that is, each community has NU / (NC * NC) people. For members located in the two-dimensional space, each time step (which can be understood as daily) moves to the neighbor community with a probability of M (u, t). Therefore, for individuals in different community, there are five possible directions for movement: no movement, up, down, left, and right(Figure 2).

**Figure 2.**
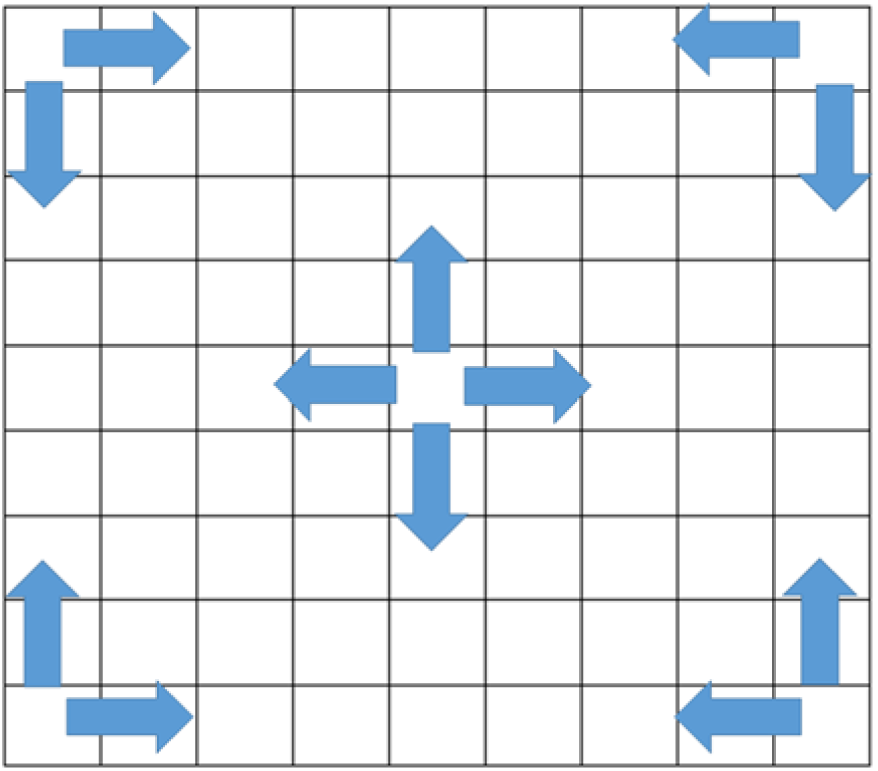
Individual move model.

## 3 Experimental design

Based on MATLAB R2017a, computational experiments were performed to simulate the epidemic prevention and control measures. Parameter settings of computational experiments are shown in Table 1.

**Table 1.**
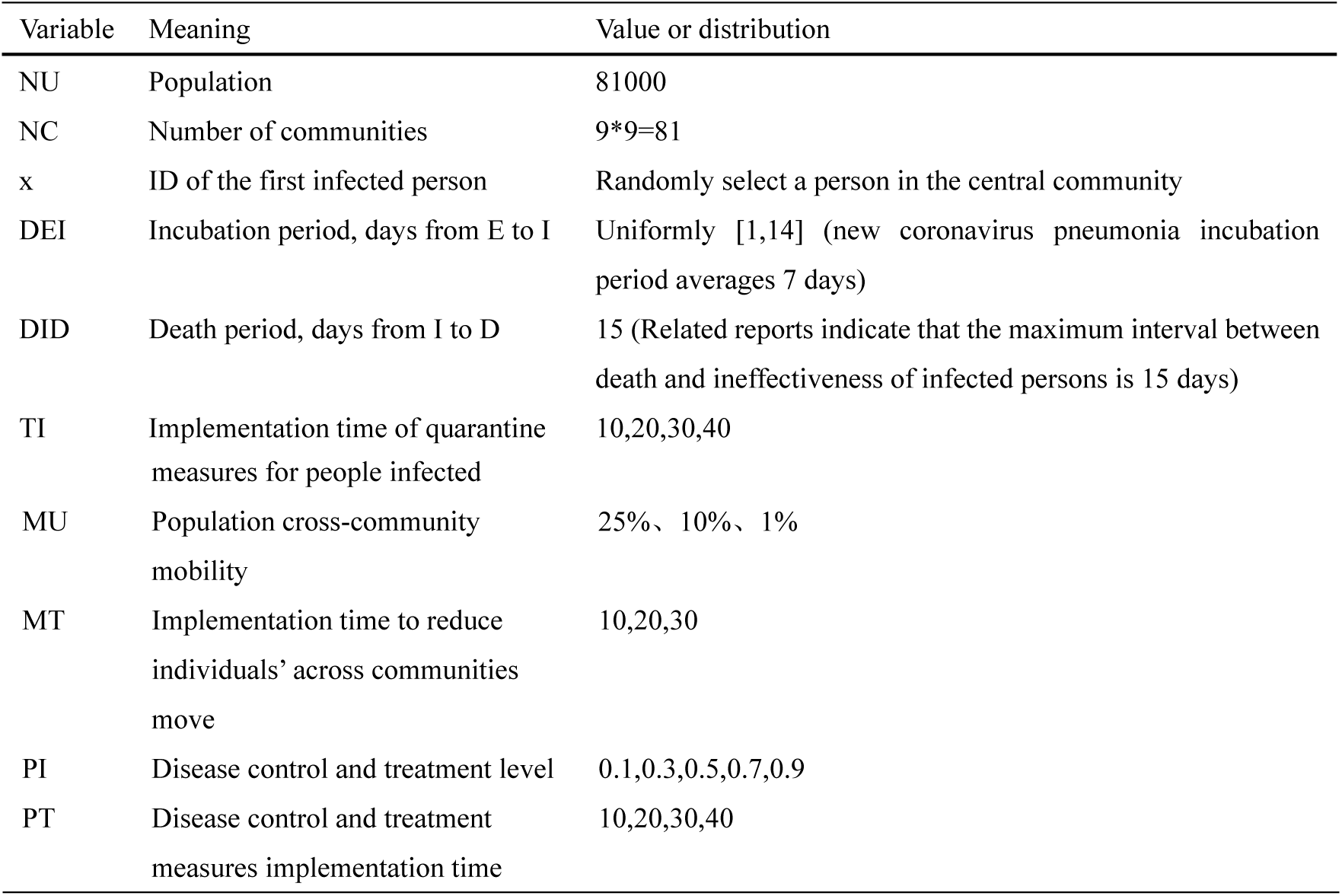
Parameter settings.

This paper conducts three experiments

- Frist, isolate the infected. In this study, the effect of isolation measures for infected people is that from the time TI, the individual in I status stopped moving and will be isolated and treated by hospital.
- Second, reduce individuals’ across communities move. In this study, measures to reduce individuals’ move is that from the time MT, reduction in the probability (MU)of all personnel going to other communities.
- Third, improve the level of treatment. In this study, the measures to improve the level of treatment is that from the time of implementation of treatment(PT), all individuals in I status will be isolated and treated with probability PI.

## 4 Simulation of infected isolation measures

In this study, the isolation measures refer to that from the time of publication TI, individuals in I-status stop moving and are isolated and treated by the hospital. Fixed relevant parameters (MU = 0.25, MT = 1, PI = 0.1, PT = 40), simulate the implementation of quarantine measures at four time points of TI = 10, 20, 30, 40 after the outbreak, and observe individual proportion changes of E and I status (Figure 3). It can be found that with the isolation measures implemented at four time points, the E-status user changes are not obvious. The possible reason is that this isolation is only implemented for I-status users, but one of the characteristics of 2019-nCoV is that not only I-status is contagious, but E-status is also contagious. Although the I-status is being treated in isolation, there are still many members of the E-status in the incubation period. These members make the disease likely to continue to spread. This explains that the government requires that people who share a bus, a flight, or a train with the infected person need to isolate themselves. These individuals may have changed to the E-status because they had been in the same space with the I-status individuals.

**Figure 3.**
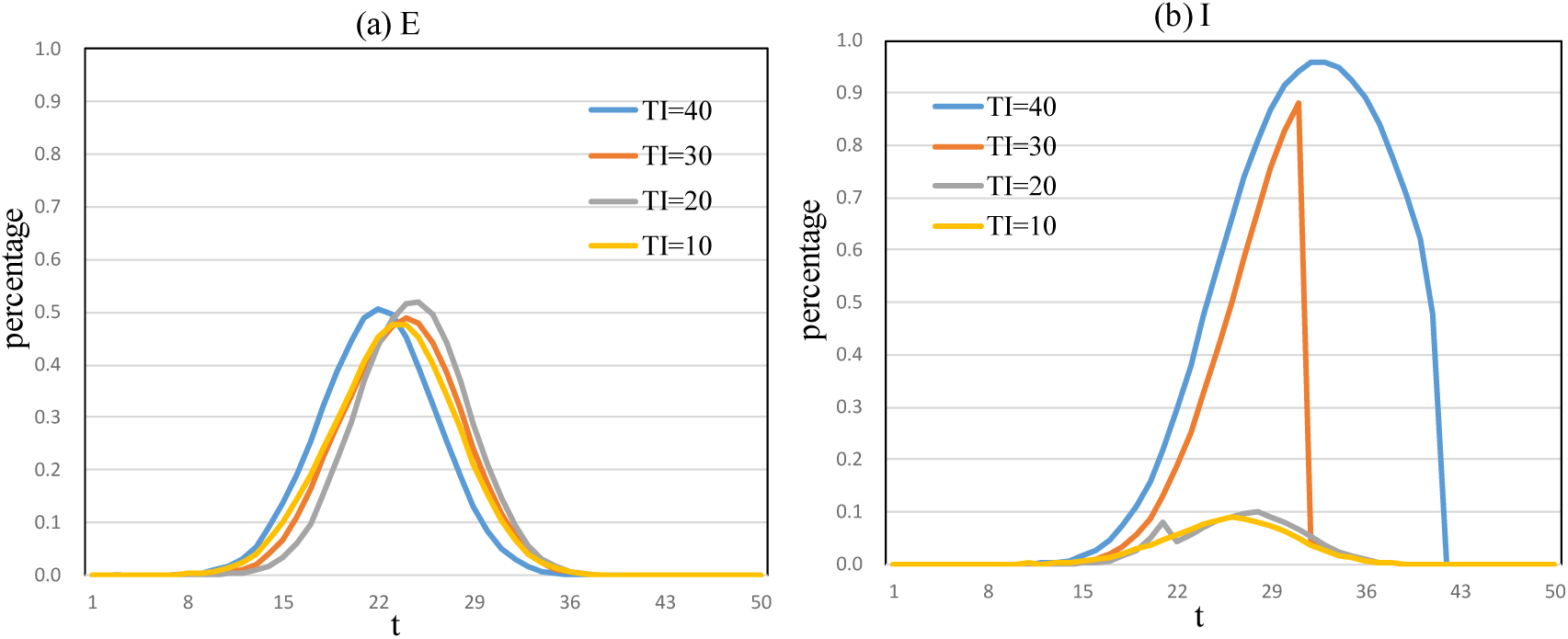
E and I-status under infected person isolation measures.

It can be found that with the implementation of isolation measures at four time points, the I-status user has changed significantly. The change is reflected in that the sooner the isolation measures are implemented, the lower the proportion of I-status users remains, and the shorter the time it takes for the proportion of I-status users to decrease to zero. For example, the peak value of I-status personnel is 10% at TI = 10, and the peak value of I-status personnel is 95% at TI = 40 and decreases to 0 at t = 45.

Individuals with infectious capacity (in E status and I status) are the key to the control of infectious diseases, but the number of deaths (in D status) is also an important manifestation of the epidemic hazard and social prevention and control capabilities. Take TI = 40 as an example, observe the proportion of D-status users in the initial community in each time period (Figure 4). At t = 20, only D-status individuals appear in the central community and surrounding neighborhoods. When t = 30, D-status individuals also appeared in a larger range of surrounding communities, but the proportion was still the highest in the central community, and the surrounding communities gradually decreased. When t = 40, the proportion of D-status individuals in the central community has reached 80%. When t = 50 (evolution into a steady state), many communities face up to 80% of D-status individuals. The epidemic evolution is centered with the central community. The larger the radius, the lower the D-status ratio. Due to the implementation of prevention and control measures, it will eventually enter a steady state, that is, there will no longer be new D-status users in all communities.

**Figure 4.**
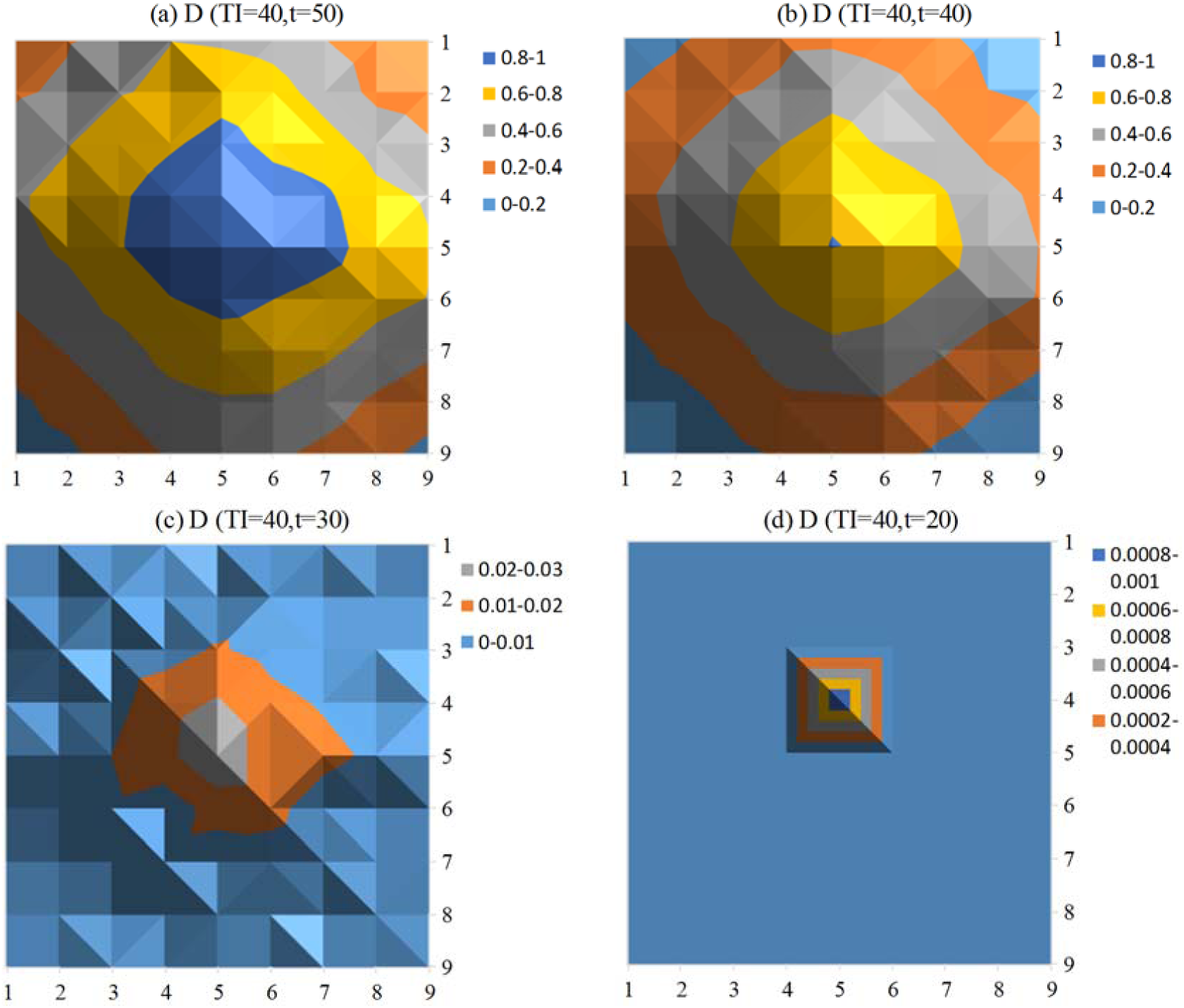
D-status under infected person isolation measures.

## 4 Simulation of reduce individuals move measures

Starting at 10:00 on January 23, 2020, Wuhan and surrounding cities announced the suspension of public transportation services to prevent the spread of new coronavirus epidemics. In this study, measures to reduce personnel mobility are reflected in the reduction in the probability (MU) of individuals going to other communities, from the time of release MT.

Fix the relevant parameters (PI = 0.1, PT = 40, TI = 200), and simulate the impact of individual mobility measures on the spread of the epidemic after the outbreak (MT = 10,20). The degree of reduction includes 0.1 (10% probability of going to neighboring communities) and 0.01 (1% probability of going to neighboring communities, similar to the city closure policy). Observe the changes in the proportion of individuals in the E and I-status (Figure 5).

**Figure 5.**
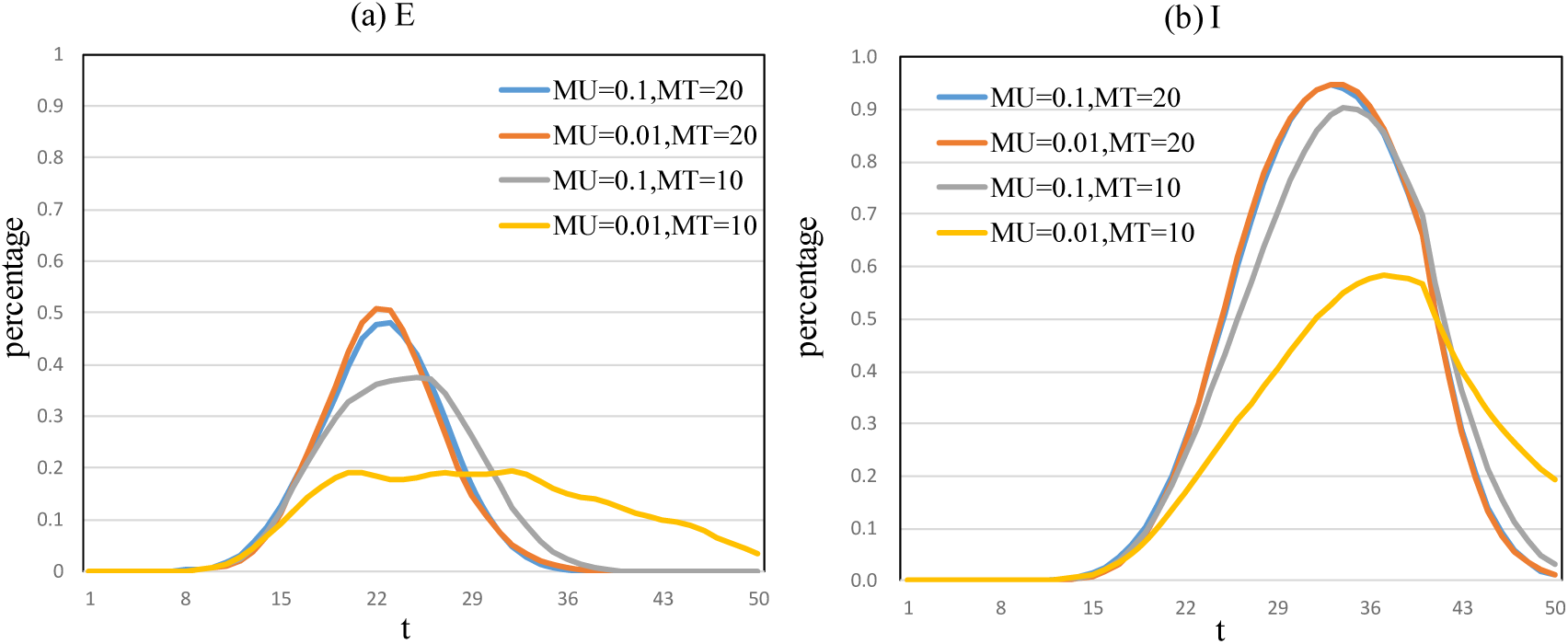
E and I-status under reduce individuals move measures.

It can be found that the implementation of population mobility control at two time points (MT = 10,20) and the different degree of population mobility control (MU = 0.1,0.01), the proportion of E-status users has changed significantly. The earlier the implementation time, the higher the degree of control, the lower the peak value of the proportion of E-status users (such as MU = 0.01, MT = 10, the peak value is 20%). In contrast, if this measure is implemented late (MT = 20), the influence of different levels of flow control measures on the proportion of E-status users is not obvious (such as MU = 0.01, 0.1, the peaks are close to 50 %). Therefore, in order to reduce the proportion of E-status users, measures to reduce population mobility should be implemented as soon as possible. If it is implemented later, even the closure of the city may be difficult to achieve significant results.

It can be found that the implementation of population mobility control at two points in time, and the difference in the degree of population mobility control, the proportion of I-status users has changed significantly. A conclusion similar to that of the E-status is obtained, that is, in order to reduce the proportion of the I-status users, measures to reduce population mobility should be implemented as soon as possible.

In order to compare the proportion of D-status users under different flow control measures, take two time stage t = 30 and t = 50 (evolving into steady state) as examples, and observe the changes in D-status users in 81 communities in the two groups (high measures,MU = 0.01, MT = 10; low measures,MU = 0.1, MT = 20) (Figure 6). At t = 30, compared with the low-measure group, the proportion of D-status in the high-measure group (MU = 0.01, MT = 10) was lower. When t = 50, compared with the low-measure group (the percentage of D-status in almost all communities exceeds 50%), the range of the D-status ratio in the high-measure group is [0.5-1]. Areas above 100% (gray) appear at t = 50, which is because the random flow of people makes the number of people in some communities higher than the initial stage. This result further validates that “the sooner, the higher the degree of implementation of the measures to reduce the movement of people, the better the effect of controlling the spread of the epidemic.”

**Figure 6.**
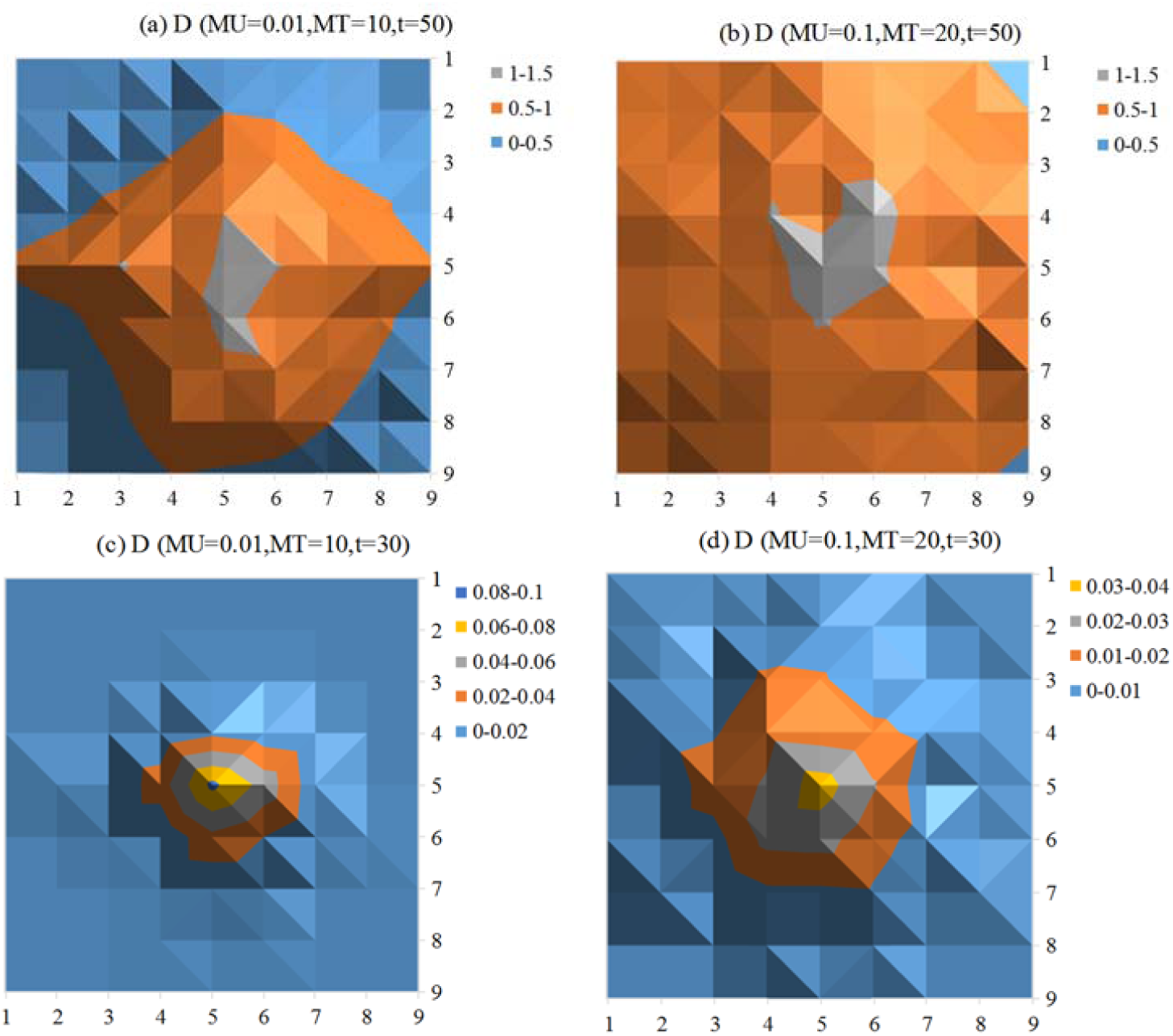
D-status under reduce individuals move measures.

## 5 Simulation of improve treatment measures

In this study, measures to improve the level of treatment means that from the time when treatment is implemented(PT), all individuals in the I state will be isolated and treated with probability PI. Fix the relevant parameters (MU = 0.25, MT = 1, TI = 200), and simulate the implementation of measures to improve treatment level at two time stages: MT = 10,20 after the outbreak. The treatment level includes 0.3 (low treatment level) and 0.7 (High level of treatment), observe the changes in the proportion of E and I-status (Figure 7).

**Figure 7.**
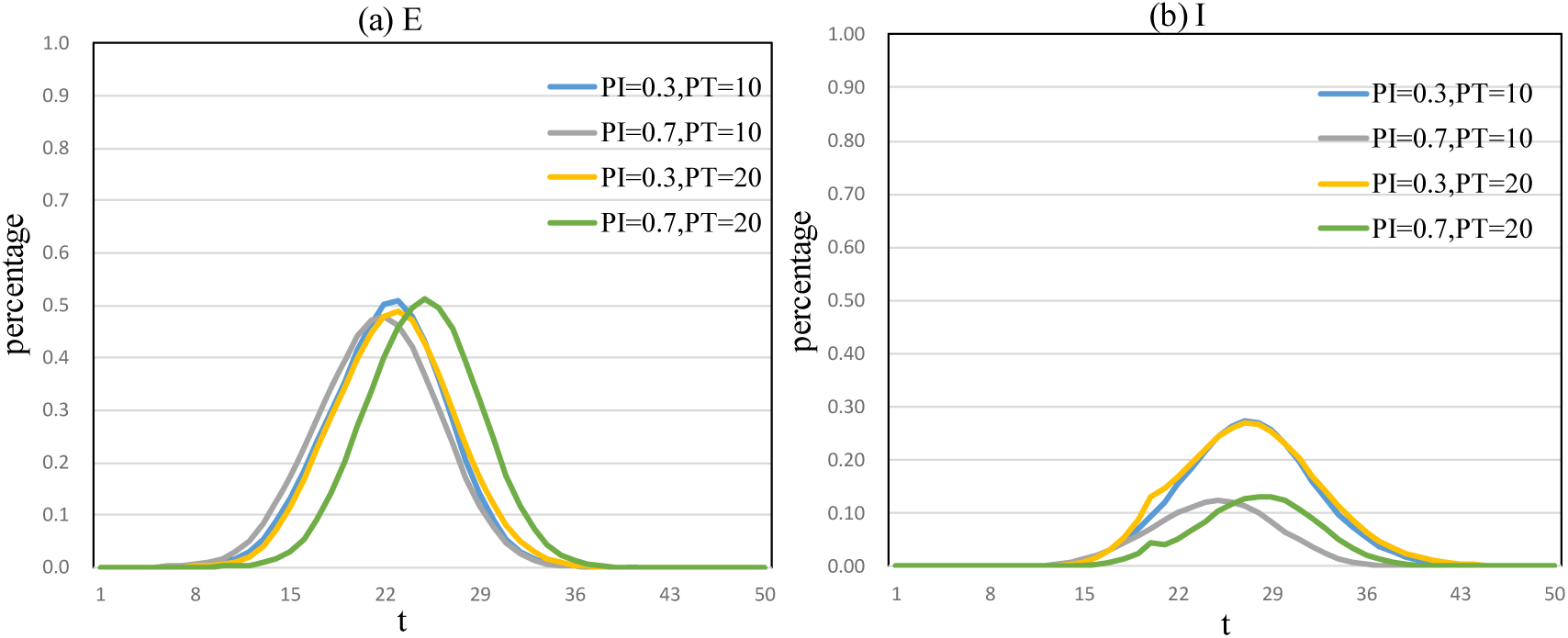
E and I-status under improve treatment measures.

From the perspective of the proportion of E-status individuals, the peak of the proportion of E-status individuals did not change significantly in the four groups. This may be because although the I-status individuals have been isolated and treated, there are still a large number of E-status individuals in the population. These individuals also have the ability to infect, that is, the ability to make other S-status individuals into E-state individuals.

From the perspective of the proportion of individuals in the I state, the peak of the proportion of individuals in the E-status changed significantly in the four groups. Among them, the group with a high level of treatment (PI = 0.7) and the early start of treatment (PT = 10) had a lower proportion of I-status individuals, and the peak appeared earlier. Therefore, starting early isolation and treatment of I-status individuals and continuously improving the level of treatment (more I-status individuals can be treated) will help reduce the proportion of I-status individuals in the population. Such findings explain why it is necessary to coordinate national medical science and technology resources to support Wuhan, and why it is necessary to build Leishenshan Hospital and Huoshenshan Hospital, because improving treatment levels as soon as possible is an important way to reduce the spread of the epidemic.

## 6 Conclusion

This study builds a SEIRD model that considers the movement of people across regions, revealing the effects of three measures (quarantine of infected people, reduction of movement of people, and improvement of treatment) on controlling the spread of the epidemic. The research results show that current prevention and control measures in China are very necessary. The results of this study are applicable to explore the impact of the implementation of relevant measures on the prevention and control of epidemic spread, and to identify key individuals that may exist during the spread of the epidemic. This study further validates the concerns of international and domestic experts regarding asymptomatic transmission (ie, E-status in this article). This article suggests that as long as medical resources are available, E-status individuals or potential E-status individuals should be included in the scope of isolation and treatment. The government should promptly release information on the epidemic situation, as well as information on the areas and vehicles used by the infected people, to further encourage those who have been in contact with individuals in the I or E-status to go to nearby hospitals for timely inspection.

## Authors’ contributions

P.S., and Y.S. designed the study. P.S. analyzed the data, carried out the analysis and performed numerical simulations. P.S. wrote the first draft of the manuscript .Y.S., and P.S. contributed to writing the paper and agreed with manuscript results and conclusions.

## References

1. Shen, M.; Peng, Z.; Xiao, Y.; Zhang, L. Modelling the epidemic trend of the 2019 novel coronavirus outbreak in China. bioRxiv 2020, 2020–2021.

2. Barthélemy, M.; Barrat, A.; Pastor-satorras, R.; Vespignani, A. Velocity and hierarchical spread of epidemic outbreaks in scale-free networks. PHYS REV LETT 2004, 92, 178701.

3. Parshani, R.; Carmi, S.; Havlin, S. Epidemic threshold for the susceptible-infectious-susceptible model on random networks. PHYS REV LETT 2010, 104, 258701.

4. Moreno, Y.; Gómez, J.B.; Pacheco, A.F. Epidemic incidence in correlated complex networks. PHYS REV E 2003, 68, 35103.

5. Kamo, M.; Sasaki, A. The effect of cross-immunity and seasonal forcing in a multi-strain epidemic model. Physica D: Nonlinear Phenomena 2002, 165, 228–241.

